# Sensitive and error-tolerant annotation of protein-coding DNA with BATH

**DOI:** 10.1101/2023.12.31.573773

**Authors:** Genevieve R. Krause, Walt Shands, Travis J. Wheeler

**Affiliations:** R. Ken Coit College of Pharmacy, University of Arizona, Tucson, Arizona, USA; Department of Computer Science, University of Montana, Missoula, Montana, USA; UC Santa Cruz Genomics Institute, Santa Cruz, California, USA

**Keywords:** Sequence Alignment, Genome Annotation, Profile Hidden Markov Models, Frameshift Mutations

## Abstract

We present BATH, a tool for highly sensitive annotation of protein-coding DNA based on direct alignment of that DNA to a database of protein sequences or profile hidden Markov models (pHMMs). BATH is built on top of the HMMER3 code base, and simplifies the annotation workflow for pHMM-based annotation by providing a straightforward input interface and easy-to-interpret output. BATH also introduces novel frameshift-aware algorithms to detect frameshift-inducing nucleotide insertions and deletions (indels). BATH matches the accuracy of HMMER3 for annotation of sequences containing no errors, and produces superior accuracy to all tested tools for annotation of sequences containing nucleotide indels. These results suggest that BATH should be used when high annotation sensitivity is required, particularly when frameshift errors are expected to interrupt protein-coding regions, as is true with long read sequencing data and in the context of pseudogenes.

## Introduction

The process of annotating the protein-coding DNA within sequenced genomes by comparison to a database of known protein sequences is called *translated search* (1, 2). Labeling of genomic sequences by comparison to proteins is generally more sensitive than comparison to the DNA that encodes those proteins (DNA-to-DNA search), due to the additional information captured in the larger amino acid alphabet and the ability to distinguish between synonymous and non-synonymous substitutions (3, 4).

A common strategy in translated search is to translate genomic open reading frames (ORFs) into putative peptides across all 6 frames, then to compare each sufficiently-long peptide to a database of previously annotated protein sequences (5, 6). We will call this approach “standard” transition. The task of translating an ORF to the encoded protein can be performed within the annotation tool (as in tblastn (7)), or by a separate preprocessing tool that may extend the ORF selection process beyond simple length filtering (such as a do novo gene prediction (8) or special handling of short ORFs (9)).

### Profile hidden Markov models

Recent years have seen tremendous gains in the speed of sequence annotation, particularly with MMseqs2 (10) and DIAMOND (11), but state of the art for maximum sensitivity continues to be observed by using profile hidden Markov models (pHMMs (12, 13)) with full Forward scores (see (10, 14) and Fig 3). These probabilistic models of sequence homology yield substantially higher sensitivity in sequence annotation (15), and serve as the basis of a wide range of annotation pipelines (10, 16– 18) and sequence-family databases (19–21). Profile HMM sensitivity gains are of paramount importance, particularly in the context of microbial community datasets, where annotation efforts often fail to identify large fractions (and in many cases, the majority) of putative proteins (22–24).

The classic tool for pHMM-based sequence annotation, HMMER3 (14), does not provide direct translated search functionality. To perform a translated search using HMMER, the user must first produce a collection of ORFs, and translate them into their encoded peptides. These candidate peptides can then be compared to the user-provided protein database using HMMER’s protein-protein search tools. Positional bookkeeping and E-value adjustment requires additional post-processing by the user.

Here, we introduce a new tool for translated pHMM search, BATH, which fills the gap left by available tools. BATH is built on top of the HMMER3 code base, and its core functionality is to provide full HMMER3 sensitivity with automatic management of 6-frame codon translation, positional bookkeeping, and E-value computation. Importantly, BATH also explicitly models nucleotide insertions or deletions (indels) that can introduce shifts in the reading frame of a protein-coding region.

### Frameshifts

A key challenge in translated search is the presence in the genomic sequence of nucleotide indels that lead to shifts in reading frame. These may represent errors during sequencing (most commonly in homopolymer regions), true mutations as in pseudogenes, or instances of programmed ribosomal frameshifting (25).

Frameshifts in protein-coding DNA lead peptide translation software using standard translation to predict fragmented peptides, which in turn may reduce the ability to identify a full-length protein, or even to annotate the protein at all. Frameshifts due to sequencing error are of concern in the context of modern long read sequencing, where indels remain a challenge (26, 27). This is particularly true for metagenomic datasets, in which low read depth for low abundance taxa (28, 29), or high population diversity as in viral genomes (30), leaves little opportunity for error correction. For example, Sheetlin et al (31) showed that the majority of metagenomic reads from a polluted soil sample contain frameshifts.

The negative effect of frameshift errors on translated alignment has been a topic of interest for over 30 years (32, 33). One way to overcome frameshift concerns is to employ a *de novo* gene finding tool that models and corrects frameshift mismatches (34, 35). An example of this approach is FragGeneScan (34), which corrects for frameshift errors while predicting ORFs in DNA sequence and is currently utilized in the MGnify pipeline (36). In principle, by correcting sequencing errors prior to ORF calls, FragGeneScan should enable annotation of some reads that are not annotated using naive ORF calls. But if error correction is overly aggressive, it can introduce new errors, causing other sequences to escape annotation. Zhang and Sun (37) showed that this concern is legitimate, and that FragGeneScan induces more errors than it fixes, leading to reduced downstream sequence homology sensitivity.

An alternative strategy is to perform homology search directly on the DNA, avoiding the calling of ORFs and explicitly modeling frameshifts. In effect, this uses homology to guide ORF prediction, which in turn leads to better homology detection. We refer to this strategy as “frameshift-aware” (FA) translated alignment. Instead of translating the target DNA into proteins and aligning amino acids to amino acids, this approach computes an alignment by comparing the amino acids of the query directly to the target DNA, assigning scores for alignment between an amino acid and a codon based on the scoring model’s value for the amino acid encoded by that codon. Under this framework, frameshifts can be addressed with the addition of what Pearson et al (38) called “quasicodons”. Rather than limiting aligned codons to the standard length of three nucleotides, Pearson’s quasi-codons could be 2 or 4 nucleotides long; other implementations (39) of this concept also include quasi-codons with lengths 1 and 5 nucleotides.

Variations on the FA approach have been implemented within both scoring-matrix based tools LAST (40) and DIAMOND (11), as well as by HMM-based tools (37, 39, 41). Of these, only DIAMOND and LAST produce E-value computations to enable evaluation of the statistical significance of their annotations, and the HMM-based approaches are all orders of magnitude slower than HMMER3. None of these methods provides full profile HMM sensitivity.

To fill this hole in the landscape of annotation software, we developed BATH. The BATH pipeline utilizes the fast filtering and accurate E-value estimates of HMMER3 to perform a combination of standard and frameshift-aware translation that maximizes the advantages of each approach while minimizing the disadvantages. BATH’s software package includes the alignment search tool and several helper tools for creating and manipulating pHMM files and is available on GitHub at https://github.com/TravisWheelerLab/BATH.

## Methods

BATH is implemented on top of the HMMER3 code base, merging aspects of *hmmsearch* and *nhmmer* pipelines with modifications specific to translated (and frameshift-aware) search. The primary tool is *bathsearch*, which requires two inputs – a protein query and a DNA target. The DNA target is simply a sequence file. The query may be provided as a BATH-formatted pHMM file containing one or more pHMMs, either created from a file containing one or more multiple sequence alignments (MSAs) using the tool *bathbuild*, or converted from a HMMER-formatted pHMM file using the tool *bathconvert*. Alternatively, the user can simply supply a file containing MSAs or independent sequences as the query input to bathsearch, which will then build the requisite model on the fly.

BATH then implements a HMMER3-like accelerated analysis pipeline (14):

- Identify ORFs within the target DNA, and convert them to peptides using standard translation. By default, all 6 frames are considered, and peptides of length ≥ 20 are retained.
- Apply HMMER3’s MSV filter to comparisons between target peptides and the set of query proteins. This computes an un-gapped alignment between each query-target pair, and retains only those pairs with a score corresponding to a P-value of 0.02 (theoretically removing 98% of unrelated sequence pairings from further consideration).
- Apply HMMER3’s Viterbi filter to remaining peptide-to-protein pairs. This computes a maximum-scoring gapped alignment for each query-target pair, then retains only those pairs with a score corresponding to a P-value of 0.001 (theoretically removing 99.9% of unrelated sequence pairings from further consideration).
- For each unfiltered match from the Viterbi stage, select a window of genomic DNA around the surviving peptide. The length *L* of the window is such that only 1e-7 of sequences emitted by the query pHMM are expected to exceed length *L* (as in *nhmmer* (17)).
- Compute a protein-to-protein Forward score, following the implementation in HMMER3’s *hmmsearch* Forward filter implementation. The score of the resulting alignment is converted to a P-value (see below), and this P-value is captured as *F*_0_.
- If *F*_0_ *>* 1*e* − 5 and *F*_1_ *>* 1*e* − 5, then filter the pair.
- For any unfiltered pair, if *F*_0_ ≤ *F*_1_, then follow a non-frameshifted (HMMER3-style) post-processing pipeline to compute protein-protein alignment boundaries, alignments, E-values, etc. Otherwise, follow an FA variant of each post-processing step, in which amino acid positions of the query are aligned to (quasi-)codons in the target.
- Compute E-values, produce translated search output that provides necessary bookkeeping information.

### Frameshift-aware model

The protein pHMM shown in Figure 1A is based on the model used in HMMER3 (14). A protein pHMM consists of transition probabilities between states (shown as arrows), emission probabilities for each amino acid within the Match (M) states (single green circles), and emission probabilities for each amino acid in the Insert states (a single grey circle). To understand how bathsearch converts a protein pHMM to an FA codon pHMM it is helpful to consider the construction of an intermediate non-FA codon model 1B. The emission probability for a codon in a match state is set to that of its encoded amino acid, under a given codon translation table. This effectively collapses all synonymous codons down to a single class, yielding probabilities that are equivalent to the protein pHMM. In the codon model, the Insert (I) states are altered so that they insert three nucleotides instead of one amino acid, but the transition probabilities that govern indel likelihood do not change.

**Fig. 1.**
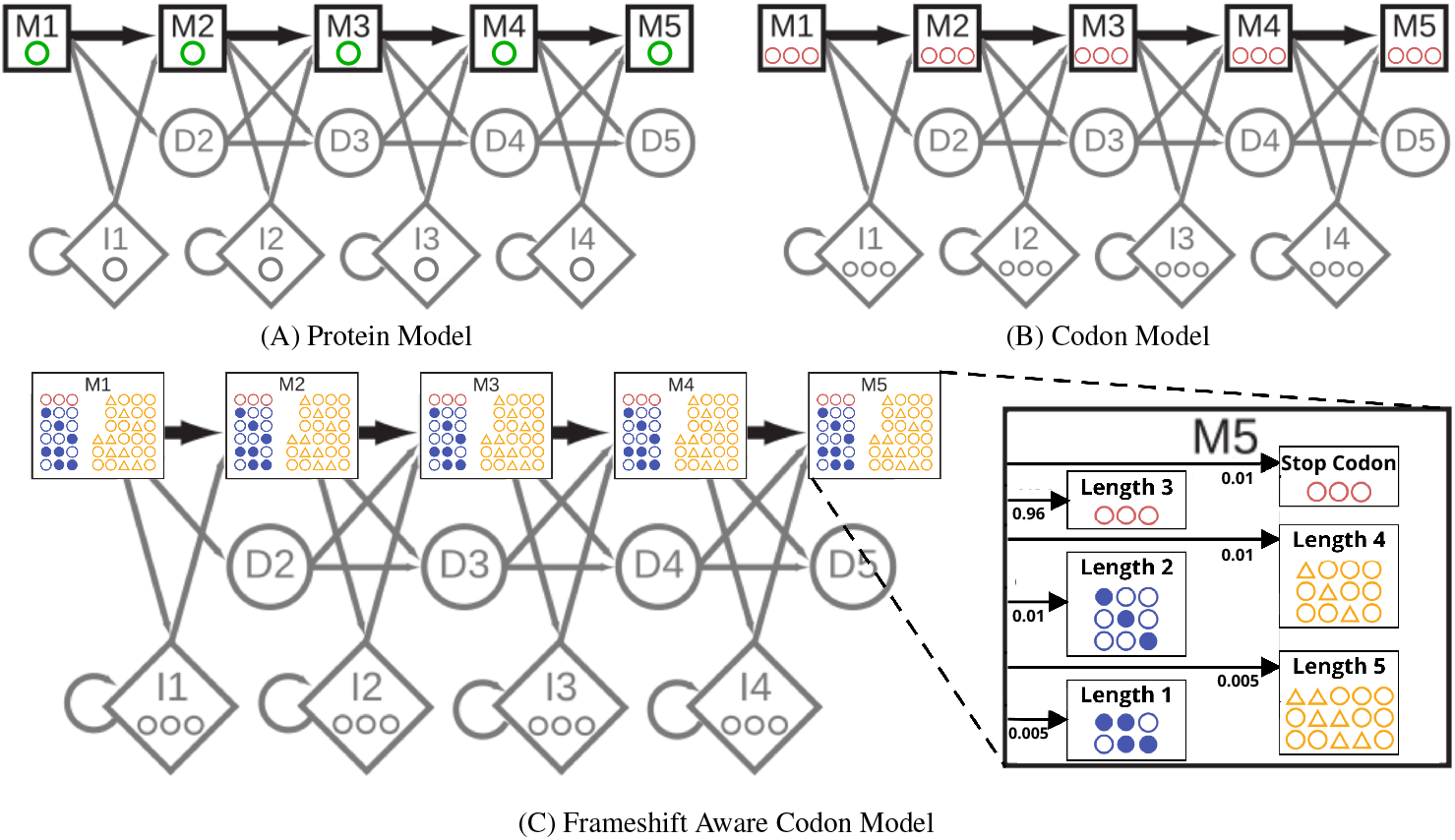
Comparison between the core model states of (A) a protein pHMM with each match state emitting a single amino acid (green), (B) a non-frameshift aware (non-FA) codon pHMM with each match state emitting a 3-nucleotide codon (red), and (C) a frameshift aware (FA) codon pHMM with each match state emitting a quasi-codon with length ranging from 1 to 5 (yellow). The protein model (A) is representative of the one used in HMMER3. The (non-FA) codon model is only an intermediary step in converting the protein model into the FA codon model in (C) which is the pHMM used by bathsearch’s FA algorithms.

To convert the codon pHMM to a FA codon pHMM (Figure 1C) additional emission probabilities need to be determined for both quasi-codons and stop codons. To identify stop codons that may arise as the result of nucleotide substitutions, bathsearch precomputes a per-state probability of emitting each stop codon *s*: for match state M_*i*_, and for each *s*, the codon *c* with the highest probability at M_*i*_ is identified among those that are at most one substitution from *s*, and the probability of *s* at M_*i*_ is set to that of *c*, multiplied by a stop-codon penalty (default 0.01).

Bathsearch allows quasi-codons of length 1, 2, 4, & 5 and assigns their emission probabilities using a similar procedure to the one it uses for stop codons. For each possible quasi-codon *q*, bathsearch considers the full set of codons *C* that could be made from *q* by either removing the extra nucleotides or adding back in the missing ones. From this set, bathsearch assigns *q* to the codon *c* ∈ *C* (and to its amino acid translation *a*) with the highest emission probability at each M_*i*_ state, and sets the emission probability of *q* to that of its assigned codon (which is the same as that of *a* from the original protein pHMM) multiplied by a frameshift penalty (default 0.01 for length 2 or 4 quasi-codons and 0.005 for length 1 or 5 quasi-codons). The full set of emissions probabilities is then normalized (all codon emissions probabilities (except for stop codon) are multiplied by 1 minus the sum of all stop codon and quasi-codon penalties.

For both stop codons and quasi-codons, the final emissions probabilities are dominated by the penalties, but small differences caused by the mapped amino acid probabilities support reasonable choices of frameshift placement in case of ambiguity. The penalties, and the normalization of the non-penalized codons, can also be thought of as another set of transition probabilities inside each match state, as illustrated in the magnified M5 state of Figure 1C. These transition probabilities serve to model the more random distribution of mutations that arise from sequencing errors or relaxed selective pressures, whereas the amino acid derived emissions probabilities and the unchanged transition probabilities between the states maintain the specific distribution of mutations seen within the query family and among related protein sequences at large. The FA codon model also differs from the HMMER3 protein model in the so-called “special states” outside the core model, see Supplementary Figure S1 for details.

### Avoiding redundant computation

Figure 2 shows a simplified representation of the alignment paths available to a protein alignment model and the FA codon model. The corresponding dynamic programming recurrence for the FA Forward recurrence contains more than 3.6x the calculations of the standard protein Forward recurrence due to the additional transition options and large number of new potential emissions. Straightforward implementation leads to excessive memory requirements and run time, as reported with the heuristic implementation in (37). This is also true of algorithms downstream in the pipeline (e.g. Backward and posterior decoding). BATH reduces the impact of this extra complexity by only using the full set of FA algorithms on target-query pairs that (according to the P-value comparison) are more probable under an FA model of homology. Even so, straightforward implementation of the full FA model is slow. Fortunately, many of the expensive calculations are repeated identically in consecutive passes through the recurrence, so that it is possible to amortize the cost of the calculation by memoizing values after their first calculation, and re-using those values until they are no longer needed (see pseudocode in Supplementary Figure S2). This reduces the number of calculations to just 1.6x that of the standard protein pHMM Forward recurrence.

**Fig. 2.**
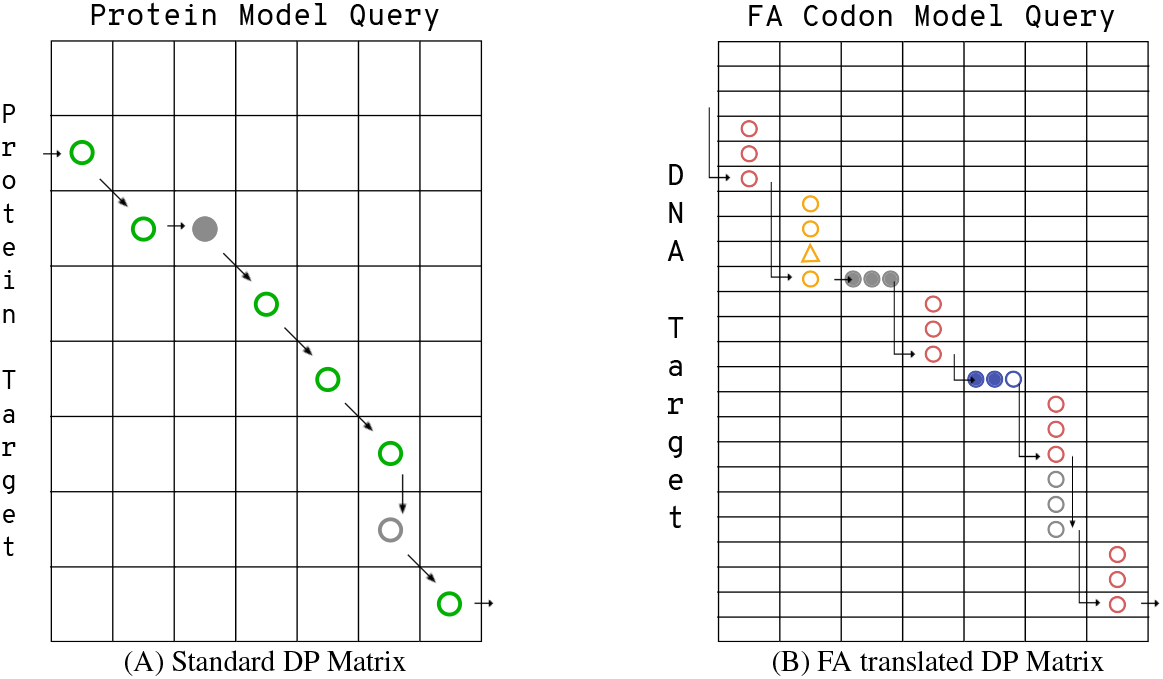
Comparison of toy DP matrices for (A) a protein alignment and (B) a frameshift-aware (FA) codon alignment. Profile HMM alignments are made up of pairings between states in the query model and residues in the target. For a protein-to-protein alignment (A), there are only three types of pairings: a match state to an amino acid (shown as a green circle) a delete state to a gap (filled grey circle), or an insert state to an amino acid (open grey circle). A codon alignment (B) has an analogous set of pairings: a match state to a codon (shown as three pink circles), a delete state to a gap (three filled grey circles), or an insert state to a codon (three open gray circles). BATH’s FA alignment also includes pairings between match states and quasi-codons (yellow circles and triangles for quasi-codons with insertions or blue filled and open circles for quasi-codons with deletions).

### Post-processing and E-value calculation

Once the target-query pair has been fully analyzed, bathsearch reports a final alignment, score, and E-value for each match that passes a user-provided E-value cutoff. Each bathsearch alignment shows the original target DNA split into (quasi-)codons, the amino acid translation of those codons, and the consensus sequence of the query protein pHMM. The final score *S* is computed in a second run of the appropriate Forward algorithm after alignment boundaries have been identified (removing non-homologous prefix and suffix regions of the target). In computing the P-value of a hit, scores are assumed to follow an exponential distribution, and parameters of the exponential are fit as described in (16). In HMMER, an E-value (indicating the expected number of false positives at a threshold of score S) is computed as a Bonferroni correction of the P-value, multiplying the P-value by the number of tested pairs for a given query. When aligning to long chromosomes, there are many overlapping candidates for alignment; in HMMER3’s DNA-to-DNA search tool, nhmmer (17), the number of tested pairs is determined by first identifying value *L* such that only 1e-7 sequences emitted by the query model will exceed length *L*, then dividing the length of the target DNA (x2 if searching both strands) by L. BATH does essentially the same thing, except it multiplies the length of the target DNA an additional x3 to count each possible translation frame.

### Construction of benchmark

To evaluate BATH in the context of other tools, we developed a benchmark that is derivative of the *profmark* benchmark used to evaluate HMMER in (14). The new benchmark, which we call *transmark*, was initiated with all the protein seed alignments from Pfam v.27 (42). Each protein sequence in those alignments was mapped to the encoding DNA using the UniProtKB IDMAP-PING resource (43). As a result, transmark contains both the protein and the DNA multiple sequence alignments (MSAs) for the Pfam families. Though several dozen individual sequences could not be mapped to their source DNA (apparently due to naming error or a mismatch in notated position), DNA source sequences could be located for at least one sequence in 14,724 of the 14,831 protein families in Pfam v.27. Since most translated alignment software, including bath-search, assumes a single NCBI codon translation Table (44) for all sequences in the target database, we eliminated all the DNA sequences that could not be correctly translated using the standard translation table (i.e. the translation did not match the Pfam protein sequence), leaving 14,569 families with at least on representative sequence. Each remaining family was then divided into a test set and a train set such that no sequence in the test set was more than 60% identical to any sequence in the train set (at the DNA level), no sequence in the test set was more than 70% identical to any other sequence in the test set, and both sets had at least ten DNA sequences. A final list of 2,673 families (202,837 test sequences and 97,497 training sequences) could be split to meet these criteria; these form the set of families and sequences used to construct a transmark benchmark.

The transmark benchmarks used here were built by selecting 1,500 of the 2,673 families at random and limiting each of those families to a maximum of 30 train sequences and 20 test sequences. The train sets were stored as MSAs (both as DNA and as proteins) to be used as queries, and the test set DNA sequences were embedded into 10 pseudo-chromosomes, of length 100MB each, to be used as targets. These pseudo-chromosomes were simulated using a 15-state HMM that was trained on real genomic sequences selected from archaeal, bacterial, and eukaryotic genomes, as in (17, 45).

In addition to 28,369 embedded test sequences, each transmark benchmark used here also embeds an additional set of 50,000 “decoy” protein-coding open reading frames into the pseudo-chromosomes. Each decoy was produced by sampling protein sequences from Pfam v.27, shuffling the amino acids to eliminate any true homology, and then using a codon lookup table to reverse translate the amino acids into DNA.

For frameshift tests, transmark benchmarks were also created with simulated indels in the embedded test sequences. Nucleotide indels were added to each test sequence at a parameterized frequency, prior to embedding into the pseudo-chromosomes. For example, at test-specific frequency of 0.02, each nucleotide position in the test sequence will serve as the starting point of an indel with 2% probability, with an equal probability of being an insertion or a deletion. An initiated indel will extend with 50%. Thus a *r* = 2% indel rate would result in an indel roughly every 17 codons, with ∼ 85.7% of those indels being not a multiple of 3 and therefore resulting in a frameshift.

## Results

To investigate the utility of BATH, we evaluated it alongside other tools for annotating protein-coding DNA. Evaluations include measurement of sensitivity when seeking true protein-coding DNA embedded within a simulated ‘genomic’ background, assessment of sensitivity in the presence of simulated frameshifts, analysis of alignment coverage and risk of alignment over-extension, and application to real bacterial sequence containing pseudogenized proteins.

### Benchmark overview

Though most evaluated tools follow a generally similar strategy involving sequence alignment of protein query to target genomic DNA, differences in implementation details lead to differences in performance, both speed and accuracy. All evaluated tools accelerate search by quickly filtering away regions deemed unlikely to contain homology, leaving only potentially homologous regions, called seeds. These seed-selection strategies may be overly restrictive, limiting the final set of matches while achieving their aim of increasing speed. Meanwhile, some of the evaluated tools are designed to produce a profile (MM-seqs2) or profile hidden Markov models (pHMMs: nhmmer, BATH) based on the available per-family multiple sequence alignments (MSAs). Profiles and pHMMs capture position-specific character frequency, and provide expected gains in sensitivity. Additionally, different frameshift models (or lack thereof) will yield different performance on sequences containing frameshifts, and other implementation details may impact the length of identified matches. We developed a series of benchmarks to explore these outcomes.

The first simulated benchmark for our analyses, *transmark00* was built as described in the Methods section as a translated search benchmark. It contains 28,369 sequences in the target set and 30,979 sequences in the query set from 1,500 Pfam families. The target DNA sequences, along with 50,000 decoy ORFs, were embedded in simulated chromosomes; insertion positions were recorded, and query sequences were retained as MSAs in both DNA and amino acid alphabets. The ‘00’ in *transmark00* alludes to the fact that the simulated frameshift insertion rate is 0% (there are no injected indels.

Some of the evaluated tools (BATH, MMseqs2, and nhmmer) are designed to produce profiles/pHMMs based on the available per-family MSAs, and then to perform sequence annotation with those profiles/pHMMs; other tools (tblastn, LAST, and DIAMOND) implement only straight-forward sequence-to-sequence annotation. To accommodate these differences we created two variants of the *transmark00* benchmark. For the first variant, each tool is given the full contingent of sequences from each family as query input; tools that can produce and use a profile or pHMM are allowed to do so, while other tools perform search with each family member in turn, adjudicating between competing matches based on best E-value. The one-search-per-sequence strategy is referred to as *family pairwise search* (46). To explore performance when per-family information is limited, a variant of the same analysis was performed using only a single query sequence per protein family (using the consensus sequence derived from the query MSAs). These variants of *transmark00*, with the full set of each family’s query sequences or with just a single consensus sequence per family, are called *transmark00-all* and *transmark00-cons* respectively.

### Tools

Analysis includes tools that are likely to be applied in current analytical pipelines for sensitive annotation of protein-coding DNA. We omit read mappers (47) as these are designed for small levels of divergence, and also ignore older tools (37, 38, 41) that are very slow, under-maintained, and are outperformed by other presented tools in our experience. For all tools, default parameters were used except where noted. E-value thresholds were set to 100 to enable analysis of tradeoffs between recall and false discovery.

We include two variants of our tool:

- *BATH* is capable of utilizing a provided protein MSA to produce a pHMM and uses that pHMM to label target DNA sequence. It may alternatively label DNA with a single protein sequence. BATH implements a frameshift model.
- *BATH --nofs* is identical to BATH, except that the frameshift model is disabled. The results from this variant are essentially the same as those produced by translating the target genomic sequence into all peptides of length ≥ 20, then running HMMER3’s *hmm-search* or *phmmer* tool (except that BATH provides positional and E-value bookkeeping).

The following tools also perform protein-to-DNA alignment. The final three tools are all faster than tblastn, and self-report BLAST-level sensitivity (at least in sensitive settings):

- *tblastn* is the relevant translated search tool from BLAST (7), and performs sequence-to-sequence comparison. This approach is not frameshift aware.
- *LAST* (31) produces frameshift-aware alignments based on sequence-to-sequence comparison. As instructed in the userguide, we used last-train to create a separate codon and frameshift scoring scheme for each benchmark.
- *DIAMOND* (11) produces frameshift-aware alignments based on sequence-to-sequence protein-to-DNA comparison. Because our primary focus is on accuracy (rather than speed), DIAMOND was run with ‘-F 15 –ultra-sensitive’.
- *MMseqs2* (10) can produce a profile from a family MSA. This enables family-to-sequence alignment with family-specific per-position scores (though note that MMseqs2 does not implement the Forward algorithm that provides much of the sensitivity seen in HMMER3 (14)). MMseqs2 is not frameshift aware.

Finally, we included one tool for DNA-to-DNA alignment, to evaluate the accuracy of searching protein-coding DNA with the DNA that encodes query proteins as opposed to the translated alignment approaches used by the other tools.

- *nhmmer* (17) performs DNA-to-DNA alignment, and can construct a pHMM from a family MSA if one is available. In general, this is expected to perform with less sensitivity than a method based on amino-acid to codon alignment, but it will also be less impacted by frameshift-inducing indels, since there is no ‘frame’ to shift at the nucleotide level. Other DNA-to-DNA tools were evaluated, but are omitted for clarity since nhmmer was the most sensitive of the group (in agreement with (17, 48)).

### Recovery of protein-coding DNA planted in simulated genomes

Using the *transmark00* benchmarks (all and cons) described above, we evaluated each tool’s ability to recover true positives while avoiding false positives. A true positive is defined as a hit that covers *>* 50% of an embedded target sequence that belongs to the same family as the query. A false positive is defined as a hit where *>* 50% of the alignment is to either an embedded decoy or the simulated chromosome. Hits are ignored when *>* 50% of the alignment is between a query from one family and an embedded test sequence from a different family, as it is not possible to discern whether true homology exists between these two instances.

Figure 3 shows that BATH (using pHMM search) identifies a larger fraction of the planted family instances, at any false positive rate, than other tools. In principle, the frameshift model of BATH could cause an increase in the score of decoys, leading to an increase in false positives; in practice, this seems not to be a concern, with the default (frameshift aware) variant producing nearly identical results to the --nofs variant. Before the first false positive, MMseqs2 (pHMM) and tblastn (family pairwise) show similar recall, but tblastn shows additional sensitivity gain with minor increases in false discovery rate. Other tools demonstrate poorer sensitivity. Despite reports of blast-like sensitivity, DIAMOND (ultrasensitive) shows much lower sensitivity in our test; surprisingly, sensitivity on this frameshift-free benchmark is improved when DIAMOND’s frameshift mode (-F) is enabled – we suspect that this is because that mode leads to slightly different parameterization of the early filter stages, which appear to be responsible for substantial reduction in sensitivity. Notably, despite ignorance of the protein-coding signal of the query, nhmmer shows sensitivity that is competitive with some of the translated alignment tools.

The results in Figure 3B for *transmark00-cons* show that BATH still outperforms the other tools even when there is only a single sequence available per query family. The relative results are similar to those seen in family pairwise search. In this single-sequence search paradigm, performance of MMseqs2 and LAST improves relative to tblastn, while nhmmer and DIAMOND see a relative degradation in performance.

The analysis shown here diverges from a recent trend of reporting recall before the first false positive (RBFFP, as in (10, 11)). We believe that reporting only RBFFP provides only partial information, as it gives no insight into the trade-off between sensitivity and false positive rates among hits in the marginal range. Furthermore, we note that the score of the first false positive is expected to grow as the number of tested decoy sequences increases. The use of large decoy sets can therefore obscure sensitivity differences in the range of moderate but important E-values. Even so, the measure is a convenient summary statistic that enables the display of the distribution of recall values across a set of queries; we present such an analysis in Supplementary Figure S3, in which BATH shows substantially greater mean and median (per query family) RBFFP. Another deviation from recent trends is that our benchmark does not make use of reversed protein sequences; this is because strings contain approximate palindromes at surprisingly high frequency, resulting in inappropriately high estimates of false discovery (49).

Our analysis is not primarily focused on speed, and the tests are not designed to explore relative performance on massive scale search, but a simple run time analysis (Table 1) shows that BATH’s frameshift model does not dramatically increase search run time relative to other methods implementing the full Forward algorithm on pHMMS (BATH -- nofs & nhmmer). The table also provides a reminder that the relative sensitivity losses of MMseqs2, LAST, and DIAMOND are offset by significant gains in speed, so that researchers primarily interested in speed should consider these as preferable options (though note that the speed of DIAMOND is under-represented in this analysis, since we used DIAMOND’s slowest settings).

**Table 1.** Run times (in hours) for all tools run on the transmark benchmark using both the family pairwise and consensus sequence query formats. All tests were run with 16 threads on a system with a 94-core Penguin Altus XE2242 @ 2.4GHz, and 512 GB RAM.

| Run Times (hours) |  |  |  |  |  |  |  |
| --- | --- | --- | --- | --- | --- | --- | --- |
| Query type | BATH | BATH (--nofs) | tblastn | MMseqs2 | nhmmer | LAST | DIAMOND |
| family pairwise | 4.34 | 1.69 | 2.31 | 0.05 | 2.20 | 0.04 | 0.22 |
| consensus sequence | 3.54 | 1.64 | 0.06 | 0.04 | 2.08 | 0.03 | 0.13 |

### Recovery of protein-coding DNA containing simulated frameshifts

Insertions and deletions (indels), whether due to sequencing error or true biological processes, can interrupt open reading frames in protein-coding DNA. The resulting frameshifts can reduce search sensitivity by depriving alignment tools of full-length peptide-coding sequences for alignment and annotation. To explore the impact of frameshift frequency, and the ability of BATH’s frameshift model to recover lost signal, we created three more transmark benchmarks with various rates of simulated frameshifts; *transmark01* has a 1% indel rate, *transmark02* has a 2% indel rate, and *transmark05* has a 5% indel rate. For more information on transmark indel rates see Methods. All of these benchmarks use the full set of query sequences (rather than a single consensus sequence).

Figure 4 mirrors the accuracy analysis of the previous section, for indel rates *r* ∈ {1%, 2%, 5%}. Unsurprisingly, tools with frameshift models (BATH, LAST, DIAMOND) see smaller degradation in performance than do the other translated search tools. BATH results demonstrate the advantage of combining pHMMs with a frameshift model. As expected, nhmmer shows only modest sensitivity loss, since it is only comparing sequences at the nucleotide level.

**Fig. 3.**
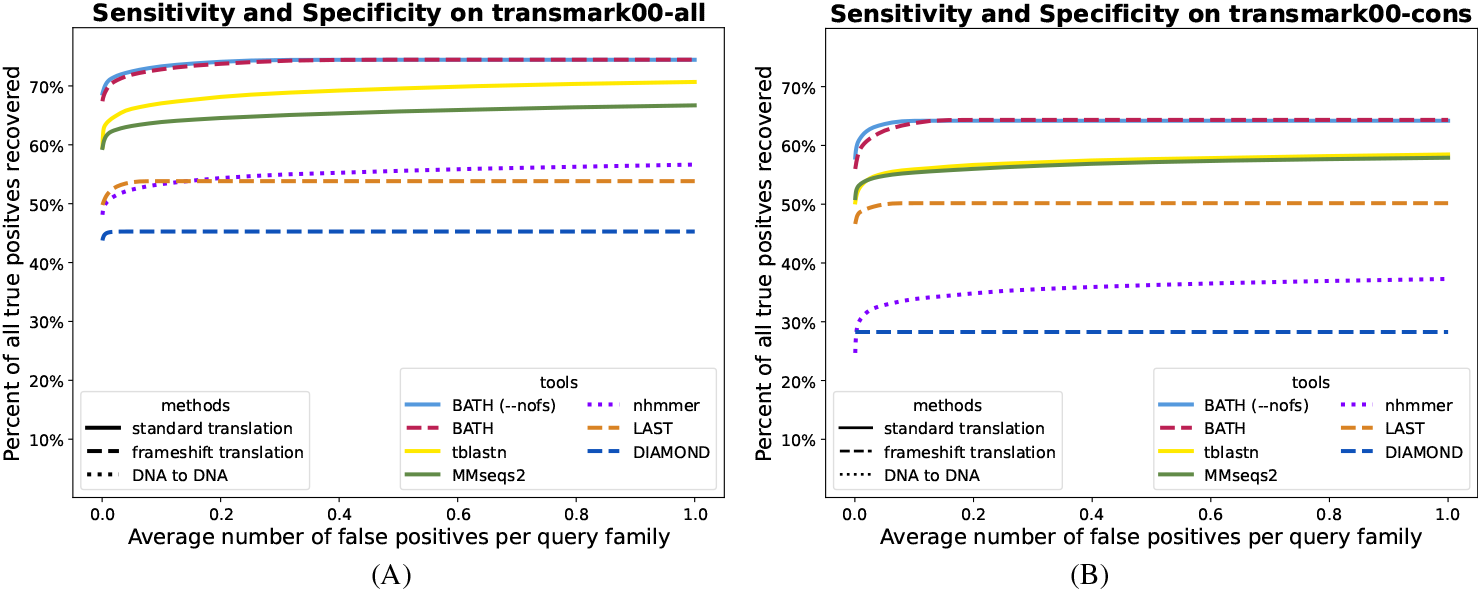
ROC plots showing sensitivity (true positives) vs specificity (false positives) for all tested tools on a translated search benchmark using either (A) 10 - 30 query sequences per family (B) a single query sequence per family. These benchmarks contain no simulated frameshifts.

**Fig. 4.**
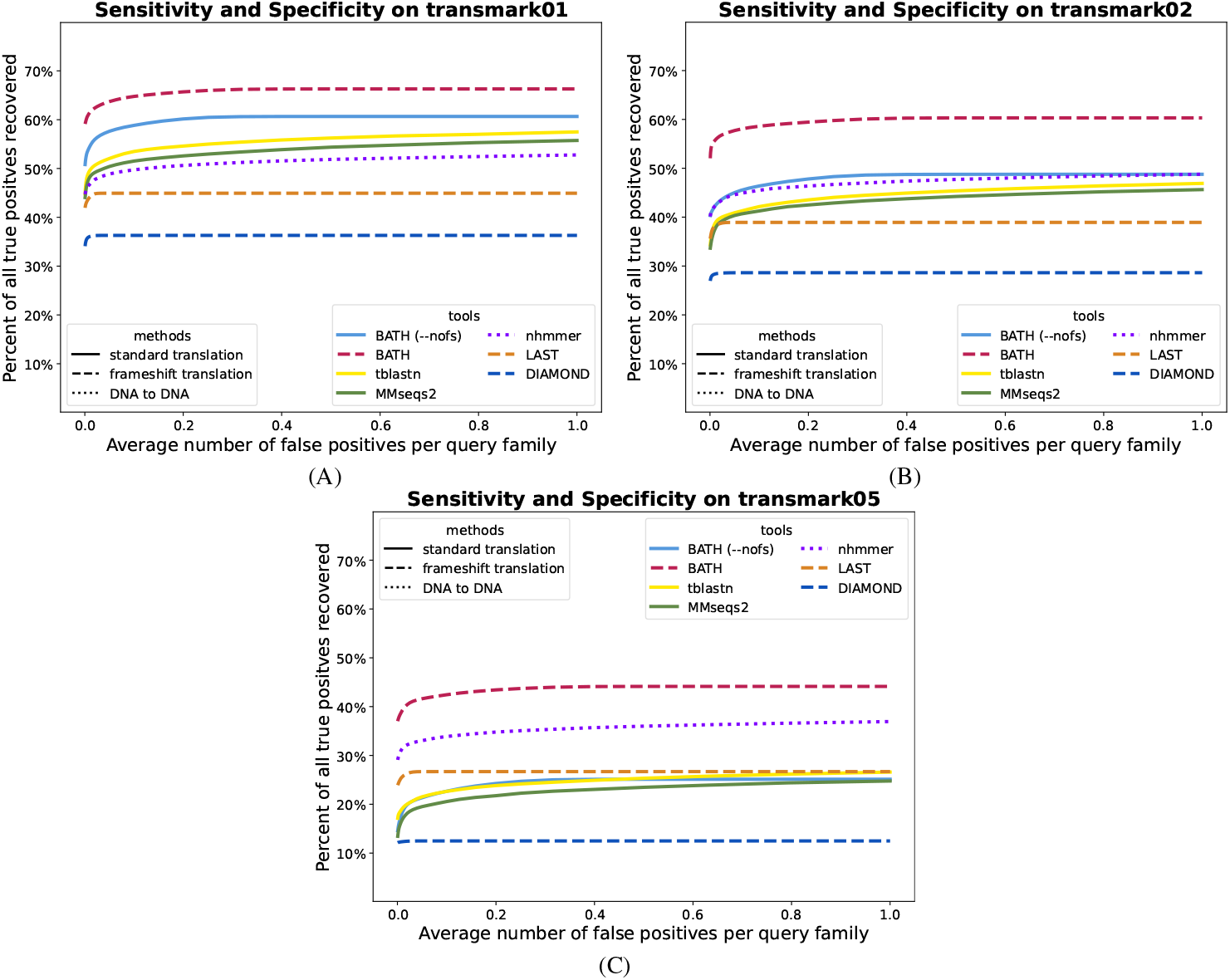
ROC plots showing sensitivity (true positives) vs specificity (false positives) for all tested tools on three translated search benchmarks with varying levels of simulated frameshift indels. (A) *transmark01* has a 1% indel rate; (B) *transmark02* has a 2% indel rate; (C) *transmark05* has a 5% indel rate.

We also analyzed the distribution across query families of recall before the first false positive (RBFFP), as shown in Supplementary Figure S3. These results indicate that some families experience total loss in sensitivity, but that in general, BATH produces substantially greater mean and median (per query family) recall than do other tools.

### Coverage and Overextension

The above assessments are measures of general recall: was a planted sequence identified, or not? Another perspective on annotation accuracy is evaluation of the extent to which the annotation accurately identifies the full length of the planted sequence – this is particularly important when target sequences are likely to be identified as fragments, as is expected for sequences containing frameshifting indels. We measure this accuracy in two ways: coverage and overextension.

To measure coverage, we identified the set of planted target sequences that were matched with E-value 1e-5 or better by all tested tools (restricting to this common set ensures that a tool is not penalized for finding a partial hit that other tools do not find at all). We computed the fraction of nucleotides in these targets that are captured in alignments produced by each tool. Table 2 presents coverage values for bench-marks contaiing 0% and 2% simulated frameshifts. In the frameshift-free test, *transmark00-all*, protein-to-DNA tools produce superior coverage to the DNA-to-DNA tool (nhmmer), and BATH’s coverage is slightly better than the others. In the frameshifted variant (*transmark02*, which also provides tools with *all* query sequences for a family), the tools with no frameshift model (tblastn, MMseqs2, BATH -- nofs) show reduced coverage due to target fragmentation. In the case of frameshifted target sequences, Table 3 shows that frameshift-aware tools (and codon-oblivious nhmmer) generally match the target sequence with a single alignment, while other tools are forced to match targets with multiple shorter fragmented matches. The summary interpretation of these results, along with those in the previous section, is that BATH find more hits (Figures 3 and 4) and also produces better coverage of the hits that it finds.

**Table 2.**
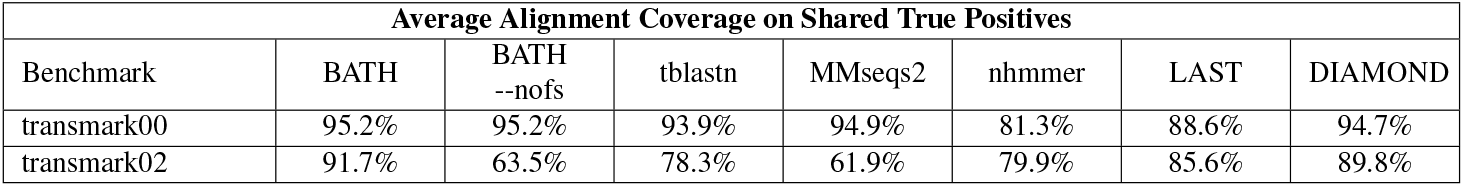
For two of the transmark benchmarks (*transmark00-all* which has no simulated frameshifts and *transmark02* which has a 2% indel rate) we calculated average alignment coverage for a set of shared true positives (the set of test sequences correctly annotated by all tools). For *transmark00-all* the set includes 10,582 hits and for *transmark02* the set includes 6,620 hits. For each sequence in these sets, we compared the true embedding coordinates of the test sequence to the alignment coordinates reported by each tool. To calculate alignment coverage, we record the percentage of all the test sequence nucleotides that are included in a true positive alignment. If there is more than one true positive alignment per tool, we combine the coverage of those alignments, being sure to only count each nucleotide once, to get the maximum coverage across all alignments.

| Average Alignment Coverage on Shared True Positives |  |  |  |  |  |  |  |
| --- | --- | --- | --- | --- | --- | --- | --- |
| Benchmark | BATH | BATH<br>--nofs | tblastn | MMseqs2 | nhmmer | LAST | DIAMOND |
| transmark00 | 95.2% | 95.2% | 93.9% | 94.9% | 81.3% | 88.6% | 94.7% |
| transmark02 | 91.7% | 63.5% | 78.3% | 61.9% | 79.9% | 85.6% | 89.8% |

**Table 3.** The average number of alignments needed for each tool to get the coverage is seen in Table 2.

| Average Number of Alignments Used to Achieve Coverage |  |  |  |  |  |  |  |
| --- | --- | --- | --- | --- | --- | --- | --- |
| Benchmark | BATH | BATH<br>--nofs | tblastn | MMseqs2 | nhmmer | LAST | DIAMOND |
| transmark00 | 1.0 | 1.0 | 1.0 | 1.0 | 1.1 | 1.0 | 1.0 |
| transmark02 | 1.0 | 2.1 | 2.1 | 2.1 | 1.1 | 1.3 | 1.0 |

Increased coverage could be the product of a simple tendency to generate long alignments. A risk created by such a tendency is that alignments may extend beyond the bounds of the true instance and into flanking non-homologous sequence, a problem known as ‘homologous overextension’ (48, 50, 51). To assess this risk, we created an overextension variant of the *transmark00-all* and *trans-mark02* benchmarks. In these variants (*transmark00-over* and *transmark02-over*), only the middle 50% of each family instance was embedded into the simulated DNA background. The result is that each full-length query match should produce an alignment to a half-length planted sequence, and any extension beyond those boundaries is a case of overextension. As in the coverage evaluation, we identified the set of planted target sequences that were matched with E-value 1e-5 or better by all tested tools, then computed measures of overextension on those hits.

Table 4 shows, for each tool, the percent of hits for which the alignment extends at least 4 nucleotides (more than a single amino acid) beyond the bounds of the true planted instance. Table 5 shows the mean length of these overextensions. Coverage and overextension are expected to correlate since a tool with low coverage will often fail to reach the end of a matched instance (incomplete coverage), and overextension is only possible after true boundaries have been reached. In line with this expectation, BATH shows a slightly elevated frequency of overextension on *transmark00-over*, though the average length of BATH’s overextensions is less than half that of the average overextension length of both tblastn and DI MOND. BATH produces longer, and slightly more frequen overextension on *transmark02-over* than on *transmark0 over*. This is due to the BATH opting to use frameshift-awa translation over standard translation far more frequently for *transmark02-over* than for *transmark00-over*.

**Table 4.** For each benchmark we first found the set of shared true positives found by all tools - 7678 hits for *transmark00-over* and 4115 hits for *transmark02-over*. From this set, we calculated the percent hits that had overextension longer than 3 nucleotides.

| Percent of Shared True Positives with Overextended Alignments longer than 3 nucleotides |  |  |  |  |  |  |  |
| --- | --- | --- | --- | --- | --- | --- | --- |
| Benchmark | BATH | BATH<br>--nofs | tblastn | MMseqs2 | nhmmer | LAST | DIAMOND |
| transmark00-over | 53.3% | 53.3% | 51.8% | 33.6% | 22.5% | 28.1% | 49.8% |
| transmark02-over | 56.8% | 28.7% | 39.0% | 18.6% | 18.7% | 38.6% | 43.0% |

**Table 5.**
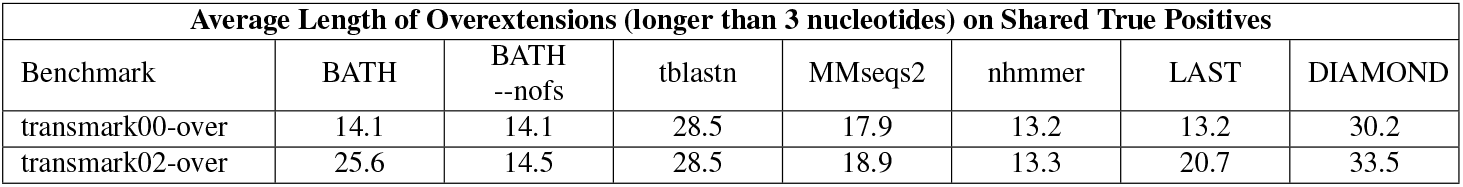
The average length of all overextensions that were at least 4 nucleotides long, taken from the set of shared true positives.

### Accuracy of E-values

Sequence annotation depends on a curate estimation of the significance of an identified matc usually provided by alignment tools with an *E-value*. For search of a given query against a given target genome *G*, pr ducing an alignment with score *S*, the E-value of an alig ment gives the number of alignments expected to meet or exceed the score *S* if *G* consisted exclusively of unrelated (random) sequences. Empirical studies demonstrate that HMMER3 E-value predictions are reasonably accurate for search against randomly generated sequences of nucleotides or amino acids (16), but these analyses do not extend to a frameshift model. To evaluate E-value reliability, we created a target set by generating 10 million random amino acid sequences of length 400 and then reverse-translated them into DNA sequences of length 1,200, ensuring each of these non-homogous decoy sequences has at least one full-length open reading frame. For the query set, we randomly selected 150 pHMMs from Pfam. We then searched these targets and queries using BATH’s three modes: --nofs which uses only standard translation, --fsonly which uses only frameshiftaware translation, and default BATH which lets the algorithm choose which form of translation to use.

Figure 5 demonstrates that estimated E-values are reasonably accurate for all three modes. The black line shows the expected occurrence of E-values less than or equal to the value of x-axis, when aligning to non-homologous sequence. The three red lines show the actual occurrence of those E-values from BATH’s three modes. The small difference seen between the lines from BATH --nofs (dotted line) and BATH --fsonly (dashed line) is explained by the impact of stop codons on the length of hits. While each of the random target sequences was generated to have at least one full-length ORF (frame 1), stop codons are often present in the other frames (frames 2 through 6). Since the standard translation used by --nofs cannot align stop codons, alignments to frames 2 to 6 will often be abbreviated compared to the same alignment from --fsonly. E-value parameterization for BATH models only the chance that a given random protein sequence will reach a target score; it does not model the chance that random DNA sequence will contain sufficiently-long ORF in all frames to produce an appropriately long random protein. When BATH is allowed to incorporate frameshifts, it overcomes the modelling challenges created by this data fragmentation, and can properly model P-values produced though both the standard and frameshift-aware Forward implementations.

**Fig. 5.**
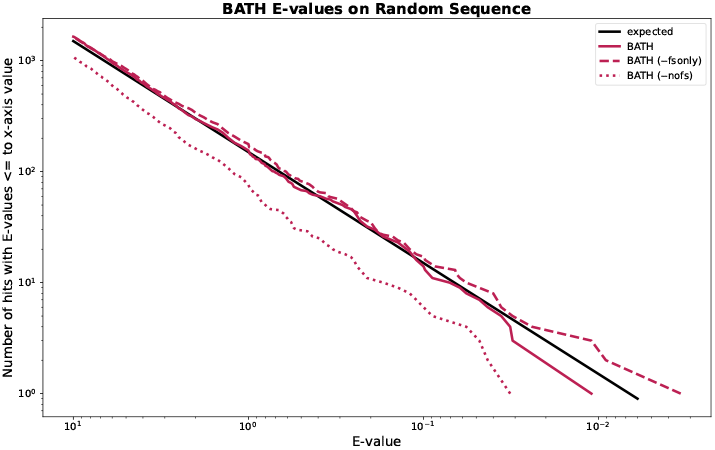
Expected and actual occurrence of E-values for BATH when running 150 randomly selected Pfam pHMMs against 10M randomly generated target sequences.

### Annotation of pseudogenes – a case study

Pseudo-genes are segments of DNA that resemble functional genes but have been rendered non-functional due to the accumulation of mutations (including frameshift inducing indels), and usually result from gene duplication or reverse transcription of an mRNA transcript.

Here, we explore the utility of BATH in annotating pseudogenes within the genomes of bacterial strains of Canditatus hodgkinia, which live as obligate endosymbionts in the cells of periodical cicadas (Magicicada) (53). The genome instability common in endosymbionts, combined with the unusual life cycle of Magicicada, has led to extreme lineage splitting among the hodgkinia in all Magicicada species. Whereas the ancestral form of Canditatus hodgkinia had only one circular chromosome, as many as 42 unique hodgkinia chromosomes have been found in a single Magicicada species. Large-scale deletions and pseudogenization have reduced genome annotation coverage from nearly 100% for single chromosome hodgkinia in other cicada species, to just 25.3% (20.2% protein-coding and 5.1% RNAs) across the hodgkinia chromosomes of Magicicadas (53, 54). The remaining sequence landscape presents a useful challenge for a tool such as BATH, since essentially all unannotated sequence is expected to be made up of pseudogenes resulting from gene loss enabled by lineage splitting and community complementation (54).

We searched 231 Canditatus hodgkinia chromosomes identified in seven Magicicada species using a curated set of 165 query protein families from hodgkinia in several other cicada species [Matt Campbell, personal comm.]. Annotation was performed with tblastn, BATH (default), and BATH --nofs. Table 6 shows the total coverage of those genomes (percent of all nucleotides involved in some annotation), produced by each tool, along with the coverage due to manual curation, as captured in GenBank. BATH (default) demonstrates clear gains in coverage, particularly in full-length hits (hits that cover at least 66% of the query length). An example of one of the Canditatus hodgkinia chromosomes (from the species Magicicada tredecim) is shown in Figure 6A, and provides examples of both novel matches (not found in GenBank or by tblastn) and improved continuity of matches that were fragmented by tblastn and BATH --nofs.

**Table 6.** Percent of nucleotides annotated by either the GenBank annotations, or by one of the three methods tested (BATH, BATH --nofs, and tblastn) across 231 Canditatus hodgkinia chromosomes from 7 Magicicada species. Only annotations with an E-value of less than 1e-5 were included in these percentages. Full-length hits are defined as hits that cover more than 66% of the length of the query. The 66% cutoff was selected based on the shortest query coverage seen in the GenBank annotations (66.67%).

| Coverage of protein-coding genes across all *Candidatus hodgekinia* chromosomes identified in *Magicicada* |  |  |  |  |
| --- | --- | --- | --- | --- |
|  | GenBank | BATH | BATH --nofs | tblastn |
| all hits | 20.2% | 37.2% | 25.8% | 30.4% |
| full-length hits | 20.2% | 33.6% | 18.6% | 19.8% |

**Fig. 6.**
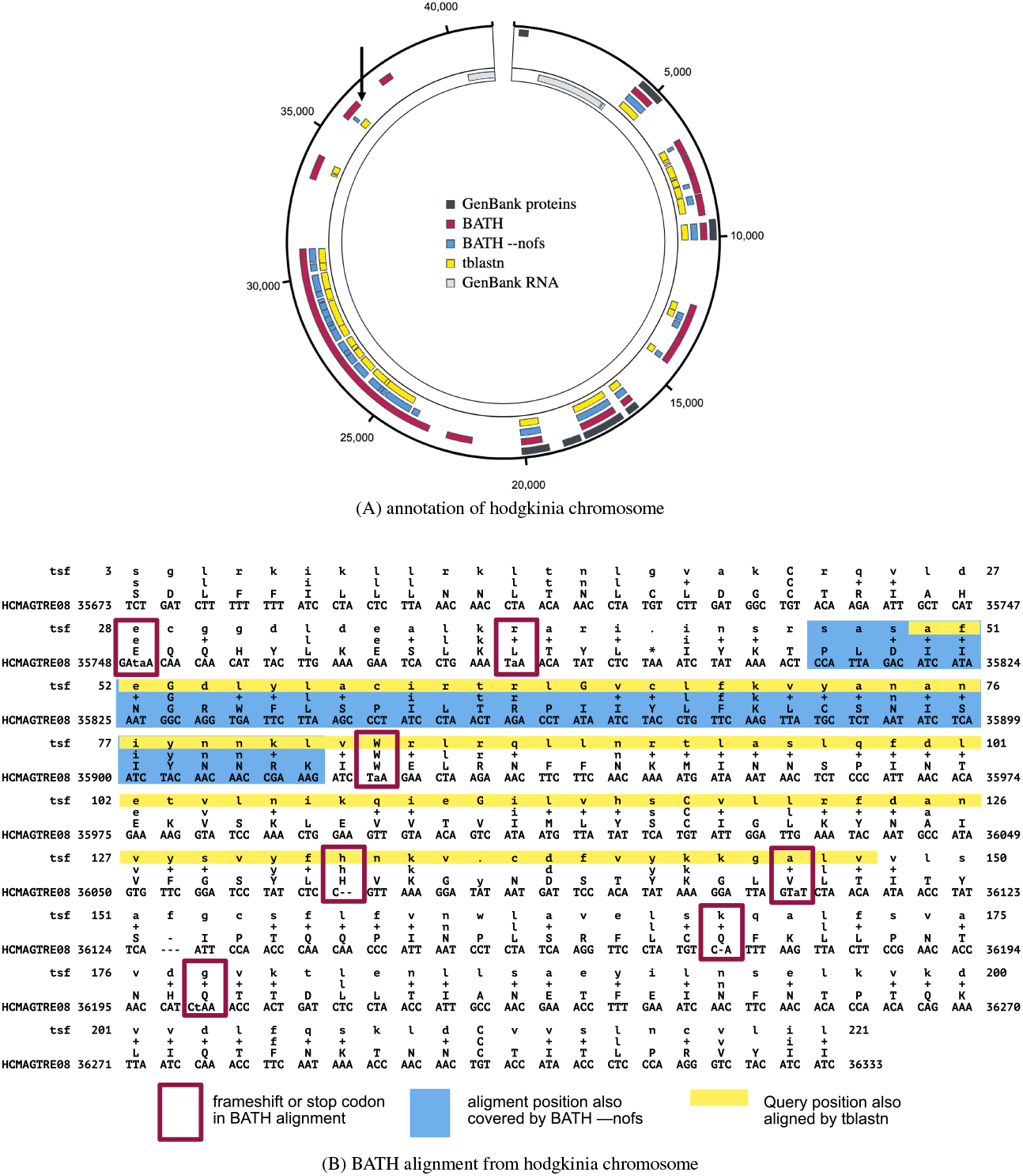
Example annotations from GenBank, BATH, BATH --nofs, and tblastn of (A) a single hodgkinia chromosome (image generated using SODA (52)) and (B) a single alignment from that chromosome. The GenBank annotations were crafted by Campbell et al. (53) with expert knowledge and a custom-built pipeline. The three tool annotations show hits with E-values less than 1e-5. The arrow in the upper left quadrant of (A) points to the location of the BATH alignment seen in (B). Each line of the BATH alignment consists of four rows. The top and bottom rows show the residues of the query (in amino acids) and target (in nucleotides), respectively. The row second from the bottom shows the amino acid translations of the target codon or quasi-codon below, and the line second from the top shows whether the match between the target and query was positive scoring (showing either the amino acid in the case of an exact match or a ‘+’ in the case of a positive scoring mismatch) or negative scoring (left blank).

Figure 6B shows a specific example of a frameshifted alignment produced by BATH, along with the abbreviated annotations from tblastn and BATH --nofs (corresponding to the annotation indicated by the arrow in the upper left section of Figure 6A). By aligning through frameshifts, BATH is able to join high-scoring regions in different frames into a single alignment. The BATH --nofs alignment is shorter than tblastn’s because tblastn’s standard translation allows for alignment to stop codons which BATH (--nofs) does not.

### False Frameshifts – a case study

One concern with the use of frameshift-aware translation is the potential that a tool will infer frameshifts that are not truly present in the sequence. We examined the risk of such “false frameshifts” by computing the rate at which BATH calls frameshifts in the true positives from *transmark00-all* and *transmark00-cons*. Of all the true positives found in *transmark00-all* fewer than 0.2% had frameshifts in their alignments and for *transmark00-cons* the occurrence of false frameshifts in true positives was just 0.4%.

Even these low false frameshift rates may overstate the true risk. The results are based on the assumption that the Pfam-sourced test sequences are correctly translated from their source DNA, but manual inspection of the highest-scoring “false frameshift” alignments from BATH all showed evidence that the protein in Pfam is translated from genomic DNA containing one or more frameshifts. One example is phosphatase CheZ from Sodalis glossinidius, a Gammaproteobacteria. The DNA that encodes this protein was aligned by BATH with a single frameshift (see Figure 7 for the alignment) not seen in the canonical translation of this protein found in Pfam. The frameshift seems plausible as it allows two high-identity regions in different frames to be stitched through a single nucleotide deletion, whereas the Pfam protein (initially sourced from Uniprot) produces a low-quality alignment in the 5’ segment. Using ESMFold (55), we predicted the structure of both the Pfam-sourced sequence and the protein sequence suggested by frameshifted alignment. Figure 8 shows a predicted disordered structure on the 5’ end of the Pfam variant that is well structured in the BATH-informed variant. Further investigation is needed to determine if this frameshift (along with others predicted by BATH during these analyses) is the result of a sequencing error or if it is a true mutation in the sequenced Sodalis glossinidius genome.

**Fig. 7.**
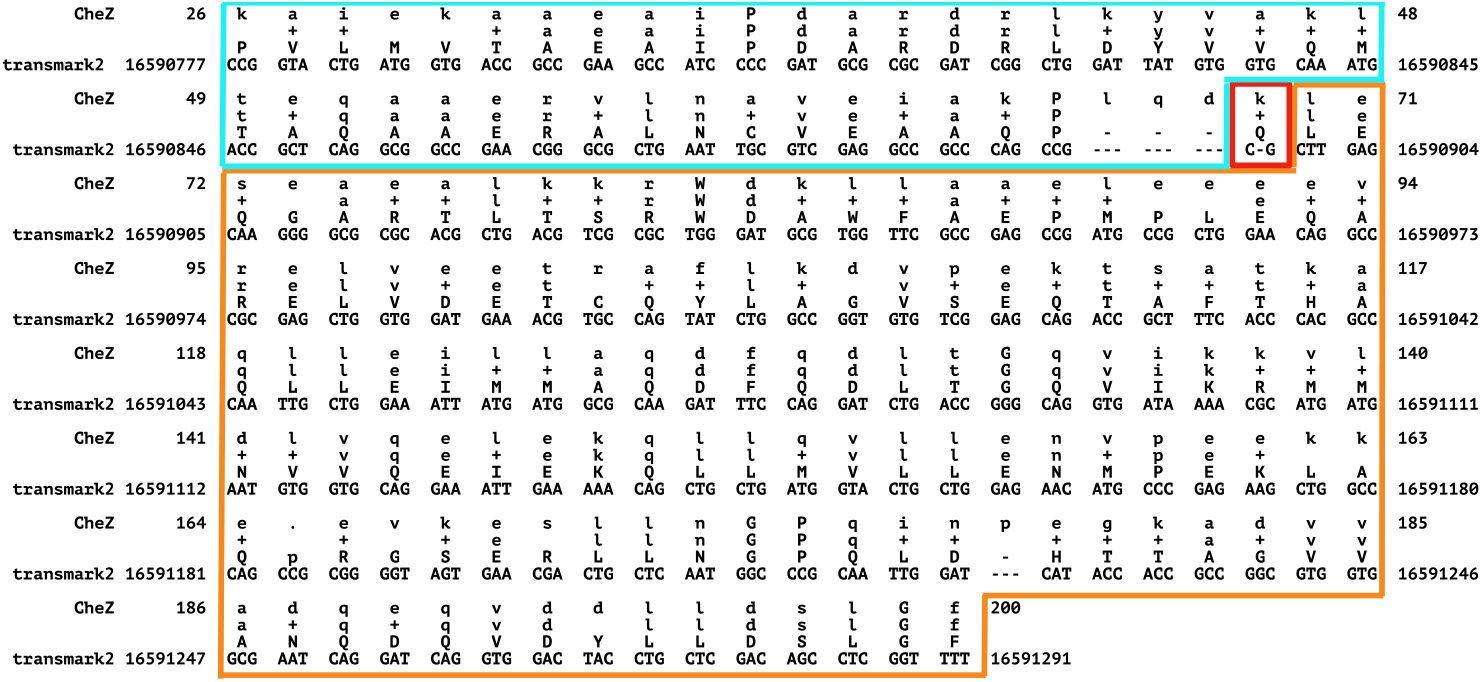
BATH alignment for the DNA encoding the CheZ protein (Uniprot:Q2NR86) from Sodalis glossinidius to the benchmark model derived from Pfam family PF04344. BATH found a single frameshift in this sequence - outlined in red. The translation after this frameshift (outlined in orange) is in the same frame as the Pfam translation, but the section before the frameshift (outlined in blue) is in a different frame and therefore has a different translation than Pfam. The BATH-predicted alignment shows high identity with the query pHMM, with 93% of the matches being positive scoring, providing strong support for the validity of the BATH translation.

**Fig. 8.**
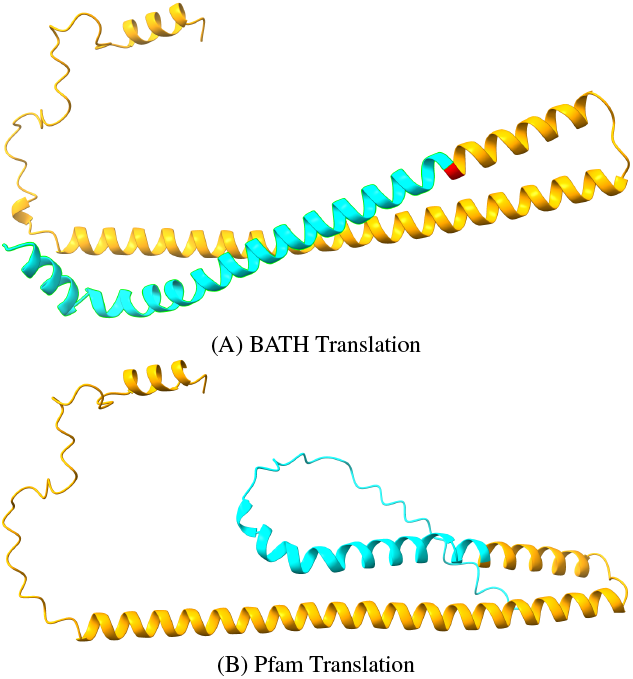
Structure Predictions for competing translations of the DNA encoding the CheZ protein (Uniprot:Q2NR86, Pfam:PF04344) from Sodalis glossinidius from (A) BATH and (B) Pfam. The orange regions show where the translations agree and the blue regions. The single red amino acid is identified as a quasi-codon by BATH, and serves as the bridge between two reading frames (see previous Figure). The blue regions differ at the level of predicted amino acid, and the Pfam translation is predicted to be disordered while the BATH translation is predicted to form an alpha helix structure.

Considering the very low rate at which BATH called any frameshifts in the *transmark00* benchmarks, combined with the fact that at least some of these frameshifts seem credible, the risk of false frameshifts from BATH appears negligible. Instead, we find that BATH’s ability to find frameshifts even where we did not expect them can lead to improved translation that could benefit existing databases.

## Conclusions

BATH provides superior sensitivity for the annotation of protein-coding DNA with or without the presence of frameshifts. It achieves this sensitivity by applying pHMMs and the Forward algorithm to the challenge of translated search and by modifying the pHMMs and alignment algorithms to be frameshift-aware. BATH also provides excellent alignment coverage and accurate E-values. Run-times are similar to other tools implementing the Forward algorithm (nhmmer) but significantly higher than other less sensitive tools (MMseqs, LAST, and DIAMOND). Future work on BATH will focus on improving run times.

## ACKNOWLEDGEMENTS

We are grateful to John McCutcheon, Matt Campbell, and Arkadiy Garber for sharing examples of clearly pseudogenized genomic sequence, discussions surrounding pseudogene and error detection, and aid in developing early versions of some annotation graphics. We thank Jack Roddy and Daniel Olson for thoughtful feed-back on both implementation and visualization. We thank Dario Copetti, Neha Sontakke, Conner Copeland, and Simon Roux for performing substantial test runs with BATH, and providing error reports and usability/documentation feedback, and also thank Sean Eddy and Nick Carter for their insightful feedback on matters of usability and formatting. We also gratefully acknowledge the computational resources and expert administration provided by the University of Montana’s Griz Shared Computing Cluster (GSCC) and the high performance computing (HPC) resources supported by the University of Arizona TRIF, UITS, and Research, Innovation, and Impact (RII) and maintained by the UArizona Research Technologies department. GK and TJW were supported during development of BATH by NIH NIGMS P20GM103546 and R01GM132600, NIH NHGRI R15HG009570 and U24HG010136, and DOE DE-SC0021216.

## Supplementary Materials

Additional supplementary materials can be found at https://github.com/TravisWheelerLab/BATH-paper, including code for construction and analysis of *trasnsmark* benchmarks.

**Fig. S1.**
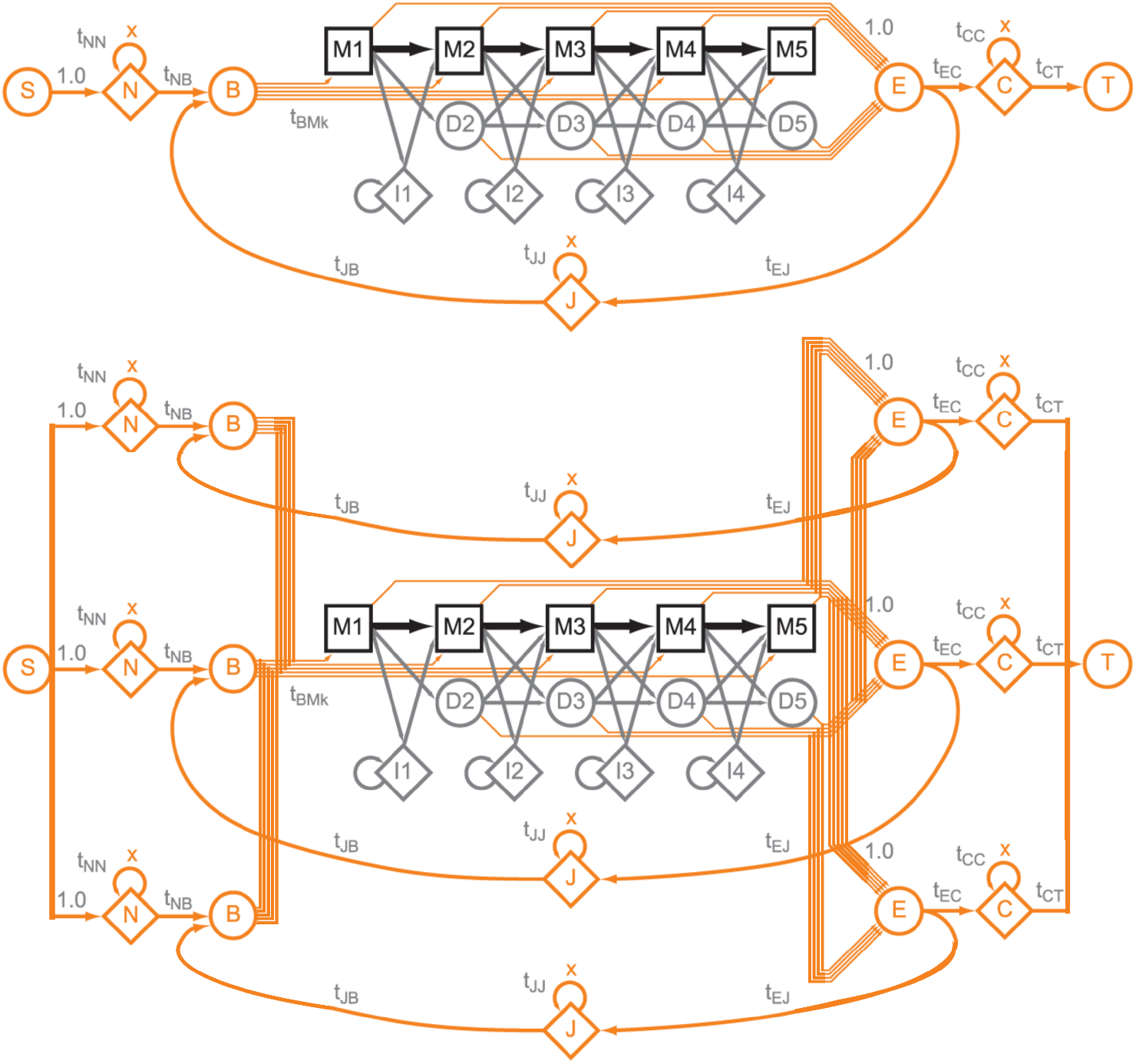
Illustration of the pHMM “special states” (shown in orange) in the protein pHMM used by HMMER3 (14) and BATH and the frameshift-aware codon pHMM used by BATH. These states allow for local alignments by permitting any non-homologous regions of the target to align to these special states rather than to the core model. The FA codon model uses three sets of the N, B, J, E, and C states, one for each translation frame. This forces the model to address any frameshifts inside the core model by emitting a quasi-codon, rather than in the special states.

**Fig. S2.**
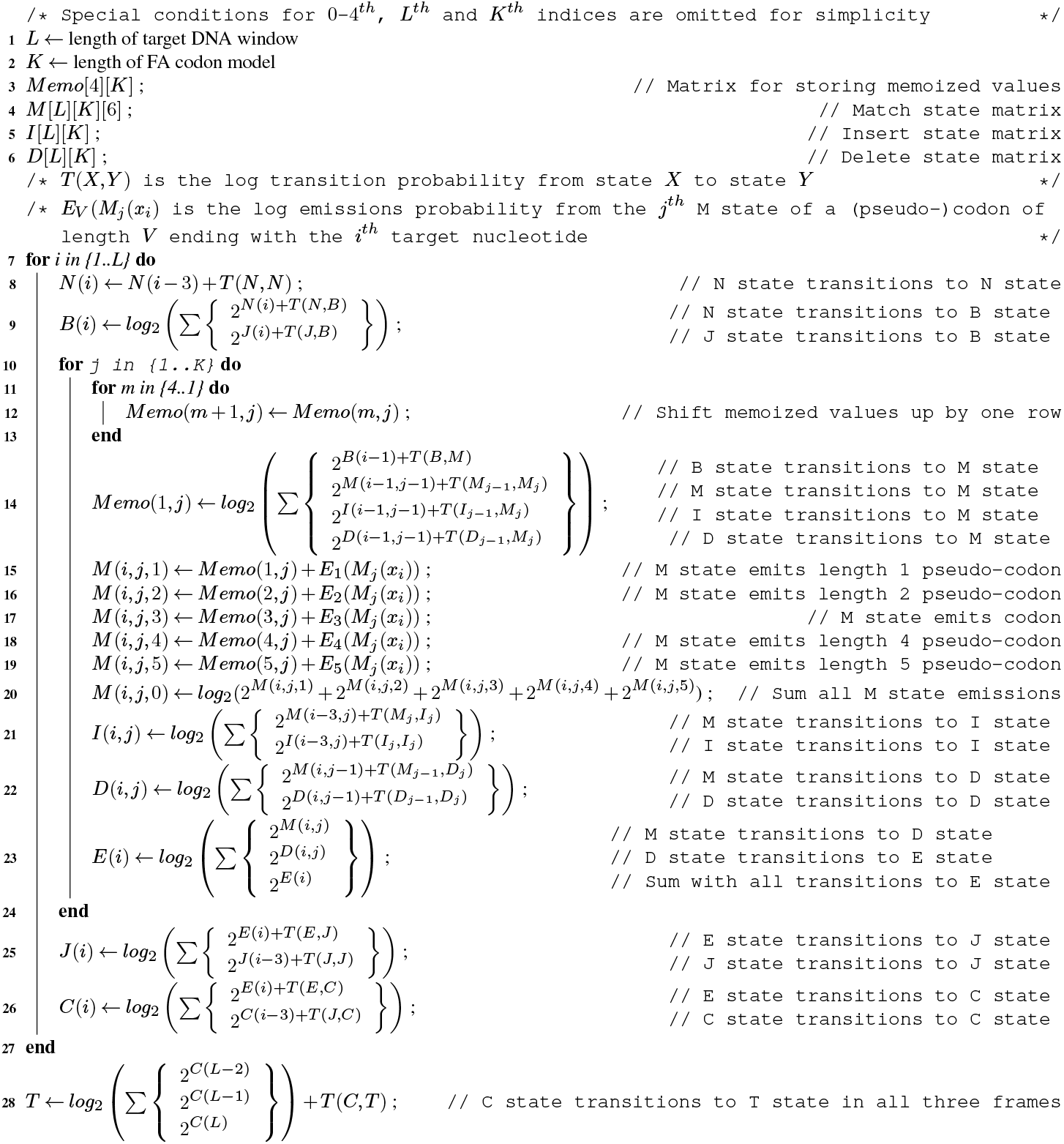
Pseudocode for the frameshift-aware Forward filter algorithm used by bathsearch. The FA algorithms employed by bathsearch use larger matrices, both because DNA targets have three times the residues of their protein counterparts and because the probabilities of various (quasi-)codon lengths often need to be stored separately, quintupling the number of M state cells. They also require more operations at each i,j position (where i is a residue in the target and j is a position in the model) to account for each of the separate (quasi-)codon emission probabilities and transition lookbacks. FA matrices also resist the application of the SIMD optimizations employed by HMMER3 for protein-to-protein alignment. In a protein-to-protein matrix such as (A), each new cell in row i can only transition from a cell in row i or row i-1. In (B), however, a cell in row i can transition from a cell in row i, i-1, 1-2, i-3, i-4, or i-5. This prevents straightforward use of the “sparse rescaling” employed in HMMER3 to prevent underflow in SIMD calculations (14). Therefore all FA algorithms in BATH are implemented without the benefit of SIMD optimization. To avoid repeated calculations, a “Memo” matrix is used to store values that will be used by subsequent M(i,j) cells. This dramatically reduces the number of additional calculations needed for frameshift-aware Forward versus standard Forward.

**Fig. S3.**
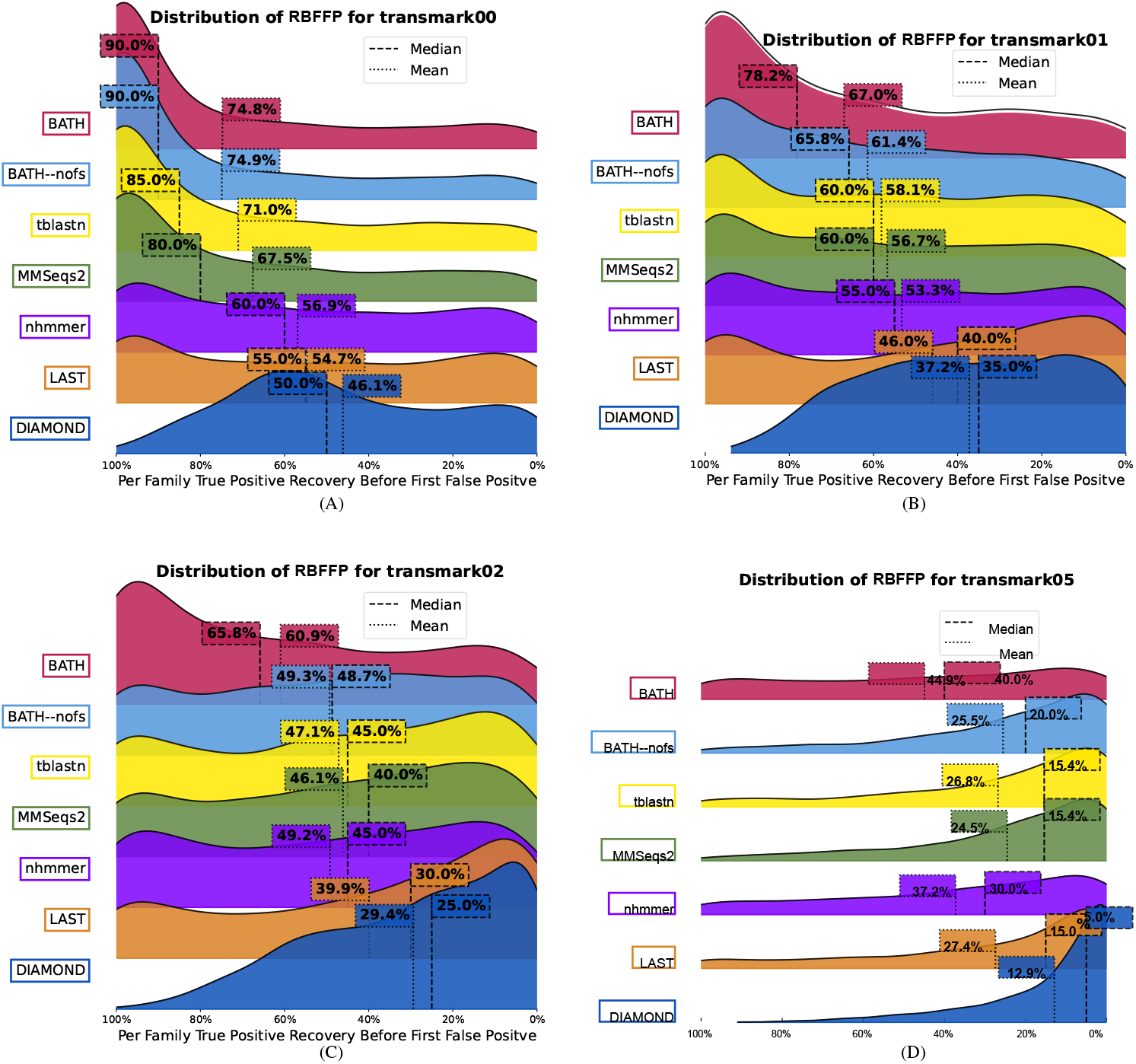
Distribution of recall before first false positive (RBFFP) values across all query family searches, with various simulated frameshift rates. In the *transmark* benchmarks, protein-coding sequences belonging to 1,500 Pfam families (up to 20 instances per family) were embedded into ten 100MB simulated genomic sequences, along with 50,000 decoy open reading frames derived from shuffled Pfam sequences. For each family, up to 30 instances are kept as a query set (see Methods for details). A true positive is defined as a hit in which *>* 50% of the alignment is between a query from a family and an embedded test sequence from the same family. A false positive is defined as a hit where *>* 50% of the alignment is to either an embedded decoy or the simulated chromosome. Hits where *>* 50% of the alignment is between a query from one family and an embedded test sequence from a different family are ignored as it is not possible to discern whether true homology exists between any two families. To find the true positive recovery before the first false positive, the true and false positives from each tool were sorted by their reported E-values (smallest to largest). If a tool had more than one hit to the same test sequence only the hit with the lowest E-value is kept. For each family, RBFFP is computed as the fraction of true (embedded) positives fpr the family that were reported with an E-value lower than the first false positive. Ridgeline plots show the distribution of these per-family RBFFP values. (A) shows that BATH produces a greater fraction of families with high RBFFP on the non-frameshifted planted sequences than do other tools. The other three plots (B-D) show the decay in mean/median RBFFP as the frequency of frameshifts increases, and that BATH experiences less decay in coverage than other translated search tools as frameshift rates increase. All tests were performed using variant in which each family is represented by the full set of training family sequences, so that the query tool can either compute a profile or perform family pairwise search.

